# Cryo-EM structure and biochemical analysis of human chemokine receptor CCR8

**DOI:** 10.1101/2023.12.30.573520

**Authors:** Qi Peng, Haihai Jiang, Xinyu Cheng, Na Wang, Sili Zhou, Yuting Zhang, Tingting Yang, Yixiang Chen, Wei Zhang, Sijia Lv, Weiwei Nan, JianFei Wang, Guo-Huang Fan, Jian Li, Jin Zhang

## Abstract

The C-C motif chemokine receptor 8 (CCR8) is a class A G-protein coupled receptor that has emerged as a promising therapeutic target in cancer and autoimmune diseases. Although the structures of human CCR8 in complex an antagonist antibody Fab1 and an endogenous agonist ligand CCL1 have been solved, the structure of ligand-free CCR8 remains to be determined. Here, we solved the cryo-electron microscopy (cryo-EM) structure of the human CCR8-G_i_ complex in the absence of a ligand. Structural analysis and comparison revealed that CCR8 in our apo structure undergoes some conformational change and is similar to that in the CCL1-CCR8 complex structure, indicating an active state. In addition, the key residues of CCR8 involved in the recognition of LMD-009, a potent nonpeptidic agonist, were investigated by mutating CCR8 and testing the calcium flux induced by LMD-009-CCR8 interaction. Two mutants of CCR8, Y113^3.32^A, and E286^7.39^A, showed a dramatically decreased ability in mediating calcium mobilization, indicating their key interaction with LMD-009. These structural and biochemical analyses provided molecular insights into the agonism and activation of CCR8 and will also facilitate CCR8-targeted therapy.

## Introduction

G protein-coupled receptors (GPCRs), located on the cell membrane, are the largest protein family encoded by the human genome [1]. Activated by chemokines or other ligands, GPCRs play an important role in transducing signals from the extracellular environment to the intracellular environment [2–4]. To date, many GPCRs have been identified in human, and they can be divided into five types according to their amino acid sequences: Rhodopsin-like receptors (Class A), Secretin receptors (Class B), Glutamate receptors (Class C), Adhesion receptors (Class D), and Frizzled receptors (Class F) [5,6]. GPCRs have been implicated in a large number of biological events, including the development of cancer and autoimmune diseases, which makes GPCRs perfect drug targets for medicinal drugs [3–7].

The C-C motif chemokine receptor 8 (CCR8) is a class A GPCR and can bind a variety of small proteins, such as CCL1, CCL8, CCL16, CCL18, with the chemokine CCL1 exclusively interacting with CCR8 [8]. CCR8 is expressed on activated T helper 2 (Th2) cells, which are an important source of pro-inflammatory cytokines, such as IL-4, IL-5, and IL-13 [9]. CCR8 is also expressed in tumor-resident regulatory T (Treg) cells [10,11]. Treg cells are known to promote cancer progression by suppressing antitumor immune responses of cytotoxic T-lymphocytes within the tumor microenvironment [12]. Depletion of Tregs in tumor microenvironment by monoclonal antibodies targeting CCR8 is currently being studied as a promising strategy in cancer immunotherapy [13,14]. Since Treg cells are endowed with immunosuppressive properties, they are also crucial in the suppression of autoimmunity [15]. It has been shown that stimulation of the CCL1-CCR8 signaling axis protected the gut from acute intestinal damage in a mouse model of inflammatory bowel disease [16]. These data indicate that CCR8 agonism is a promising strategy for the treatment of autoimmune diseases, such as multiple sclerosis, rheumatoid arthritis, and Crohn’s disease. Therefore, elucidating the structure and working mechanism of CCR8 will shed light on the development of potential intervention strategies for CCR8 related diseases. However, structural information on human CCR8 is currently limited. The structural basis for mAb1-mediated inhibition and CCL1-mediated activation of CCR8 has been reported [17], but the structures of CCR8 in apo form remain to be determined.

Various small molecule CCR8 agonists have been developed over the past decade [18–21]. Among these, LMD-009 is a nonpeptide agonist for CCR8 with high affinity in the nanomolar range, sharing the same efficacy and potency as CCL1 [20]. Thus, LMD-009 holds great potential in the treatment of autoimmune diseases. However, the molecular basis for LMD-009 mediated activation of CCR8 has not been fully elucidated.

In the present study, we employed single-particle cryo-electron microscopy (cryo-EM) to determine the structure of the human CCR8-G_i_ complex in the absence of ligand (apo) at resolutions of 2.58 Å. A detailed analysis revealed the structural characteristics of ligand-free CCR8, which displays an active state in apo structure, expanding our understanding of this drug target. Furthermore, combined with mutagenesis and calcium mobilization assay, our study revealed key residues of CCR8 involved in the LMD-009 recognition. The structural and molecular insights derived from this study will inform the CCR8 agonism by small molecules and CCR8-targeted therapy.

## Results

### Protein Expression and purification

The encoding sequence of human CCR8 was cloned into the modified pFastBac1 vector (Fig. 1A), and the recombinant plasmid was then transfected into Spodoptera frugiperda (Sf9) cells to generate baculovirus. Similarly, the dominant-negative Gα_i1_ (DNGα_i1_) [22], Gβ_1_γ_2_, and scFv16 expression plasmids were also used to generate baculoviruses. A NanoBiT tethering strategy was used for the stabilization of the CCR8-G_i_ complex [23,24]. Insect cells Sf9 co-infected with these four recombinant baculoviruses at a CCR8:DNGα_i1_:Gβ_1_γ_2_:scFv16 ratio of 1:2:1:1 to produce target protein. During the process of expression and purification, a small molecule agonist LMD-009 was added. As shown in Fig. 1B, the purified CCR8-G_i_-scFv16 complex was eluted from size exclusion chromatography (SEC) as a monomeric peak, indicating the stability and homogeneity of the target protein. The elution volume was 14.58 mL, and the molecular weight of CCR8, DNGα_i1_, Gβ_1_, Gγ_2_, and scFv16 was approximately 68, 40, 37, 8, and 30 kDa, respectively. The elution fractions were pooled and ultimately concentrated to 6.2 mg/mL for cryo-electron microscopy data collection.

**Fig. 1.**
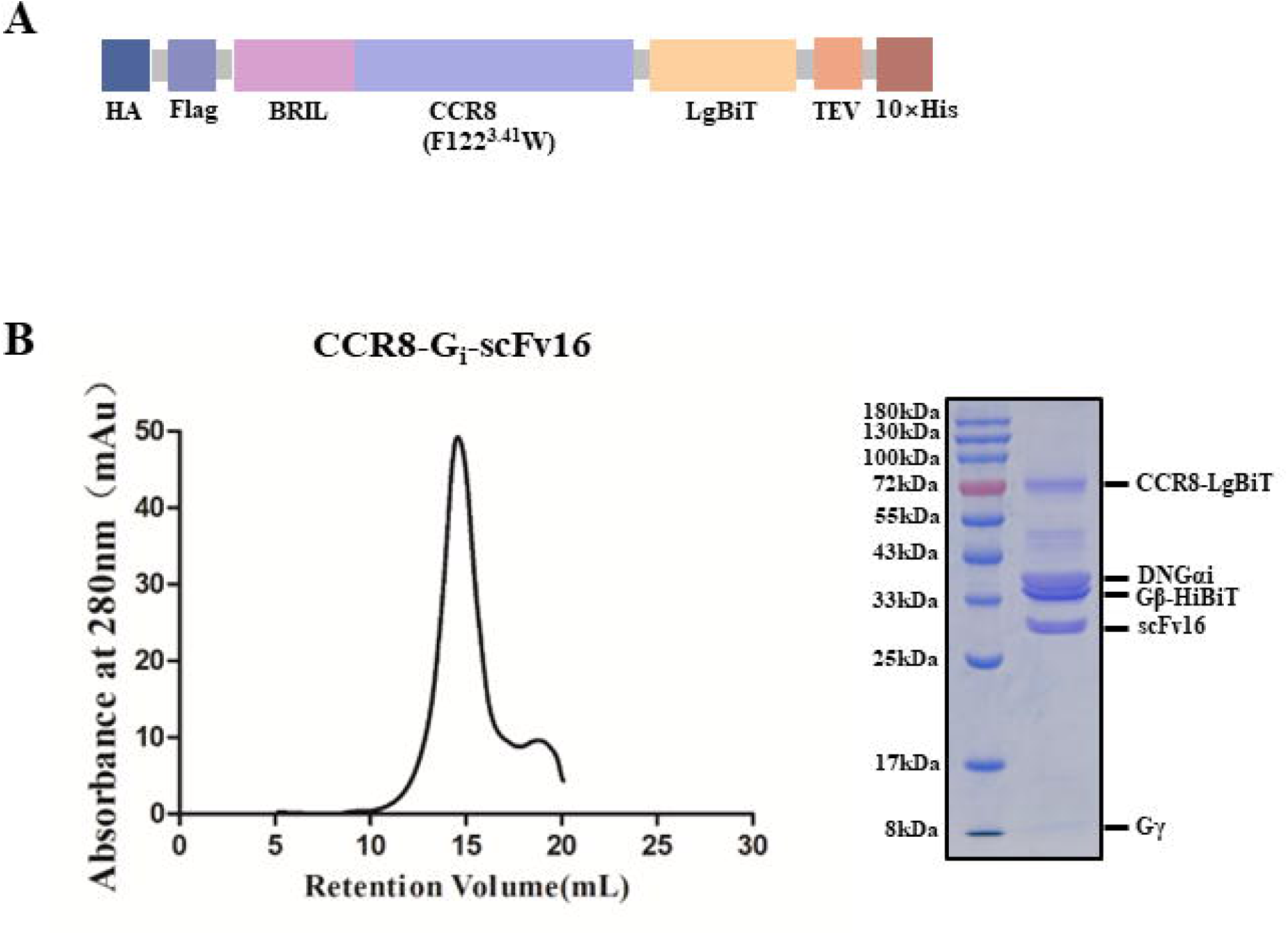
Construct, protein Expression, and purification. **(A)** A schematic diagram of the CCR8 construct is used in this study. Human CCR8 (a mutation F122^3.41^W was introduced) was cloned into a modified pFastBac1 vector, and a LgBiT was appended at the C-terminus. **(B)** Purification of the CCR8-G_i_-scFv16 complex. The recombinant plasmid was transfected into Sf9 cells. The CCR8-G_i_-scFv16 complex was expressed using an insect baculovirus expression system. The target protein was purified by using size-exclusion chromatography. The elution fractions were pooled and analyzed by SDS-PAGE.

### Overall structure of CCR8-G_i_-scFv16

The cryo-EM structure of ligand-free CCR8-G_i_-scFv16 complex was solved at 2.58 Å resolution (Fig. 2 and Fig. 3). In the CCR8-G_i_-scFv16 apo structure, the side chains of most of the amino acids in the receptor and G protein regions could be clearly defined, which enabled us to build and refine the near-atomic resolution structure (Fig. 2A and 2B). However, the flexible receptor N/C-terminus and the extracellular loops (ECL2, ECL3) of CCR8 were difficult to be observed due to their high flexibility and, therefore, could not be modeled (Fig. 3A and 3B).

**Fig. 2.**
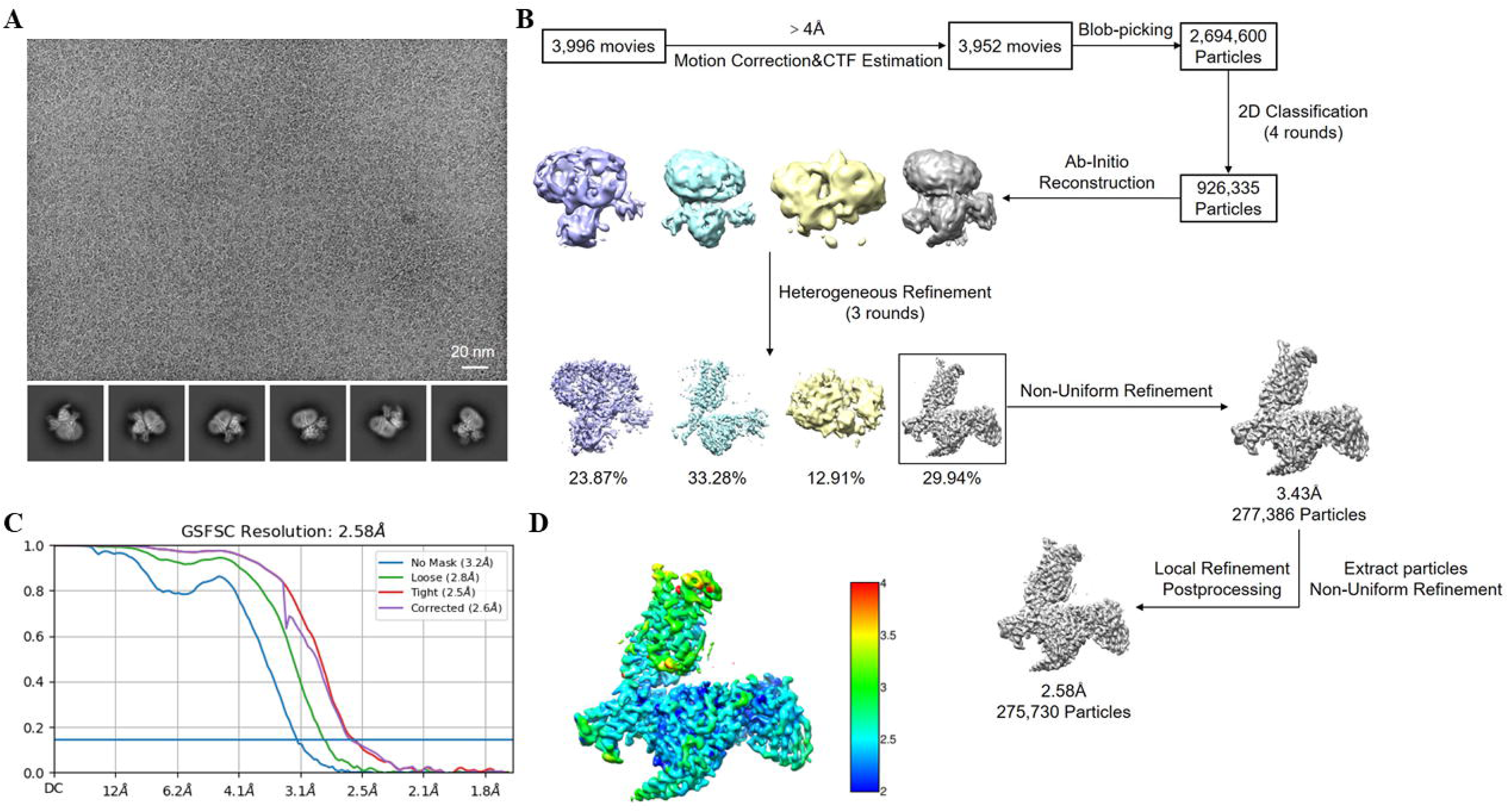
Cryo-EM data processing and structure determination. **(A)** Representative cryo-EM micrograph of CCR8-G_i_-scFv16 complex and selected two-dimensional class averages of cryo-EM particle images. **(B)** Cryo-EM data processing workflow. **(C)** Gold-standard Fourier shell correlation (FSC) curve showing an overall resolution at 2.58 Å for the CCR8-G_i_-scFv16 complex. **(D)** Cryo-EM density map colored by local resolution.

**Fig. 3.**
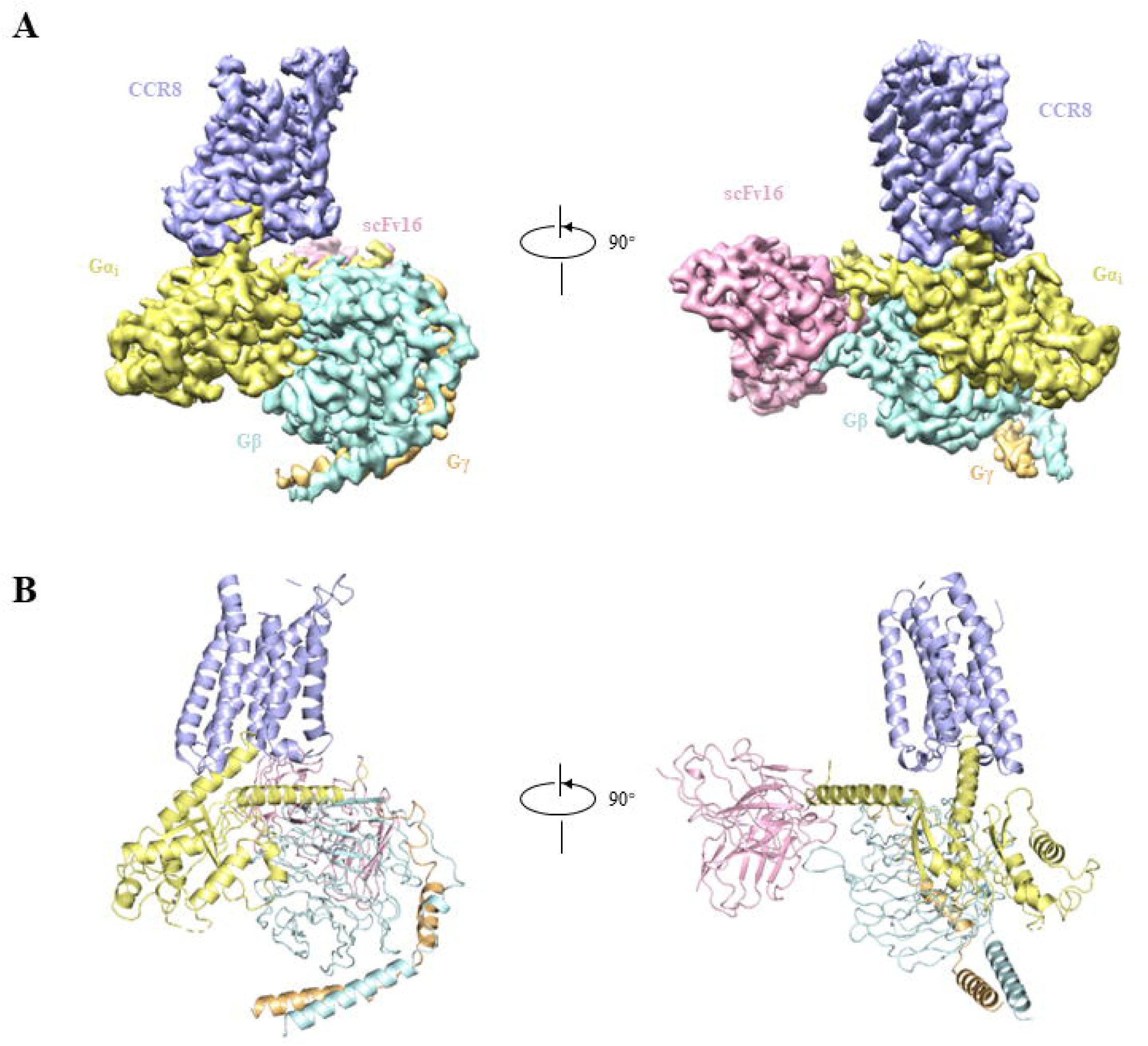
Cryo-EM structure of CCR8-G_i_-scFv16. **(A-B)** Overall structure of CCR8-G_i_-scFv16 complex. Cryo-EM density map of CCR8-G_i_-scFv16 complex **(A)** and model of CCR8-G_i_-scFv16 **(B)**, with CCR8 in light blue, Gα_i1_ in pale yellow, Gβ_1_ in pale cyan, Gγ_2_ in light orange, and scFv16 in light pink.

By comparison, we find that the overall structure of CCR8-G_i_ apo structure is similar to the structure of CCR8-G_i_ bound with an endogenous agonist ligand CCL1 (PDB ID 8U1U), with the root mean square deviation (RMSD) value of equivalent Cα positions being 0.577 Å (Fig. 4A and 4B). Comparison of CCR8-G_i_-scFv16 apo structure with the recently reported structure of CCR8-G_i_ in complex with an antagonist antibody Fab1 (PDB ID 8TLM) reveals notable structural differences, with an RMSD of 1.076 Å. Specifically, there is an inward movement of the extracellular portions of TM1, TM2, and TM5, while there is an outward movement of the intracellular side of TM6 (Fig. 4A and 4B), indicating a characteristic hallmark of CCR8 activation. The ligand LMD-009 was not seen in our refined CCR8-G_i_ complex structure, which may support the self-activation of CCR8.

**Fig. 4.**
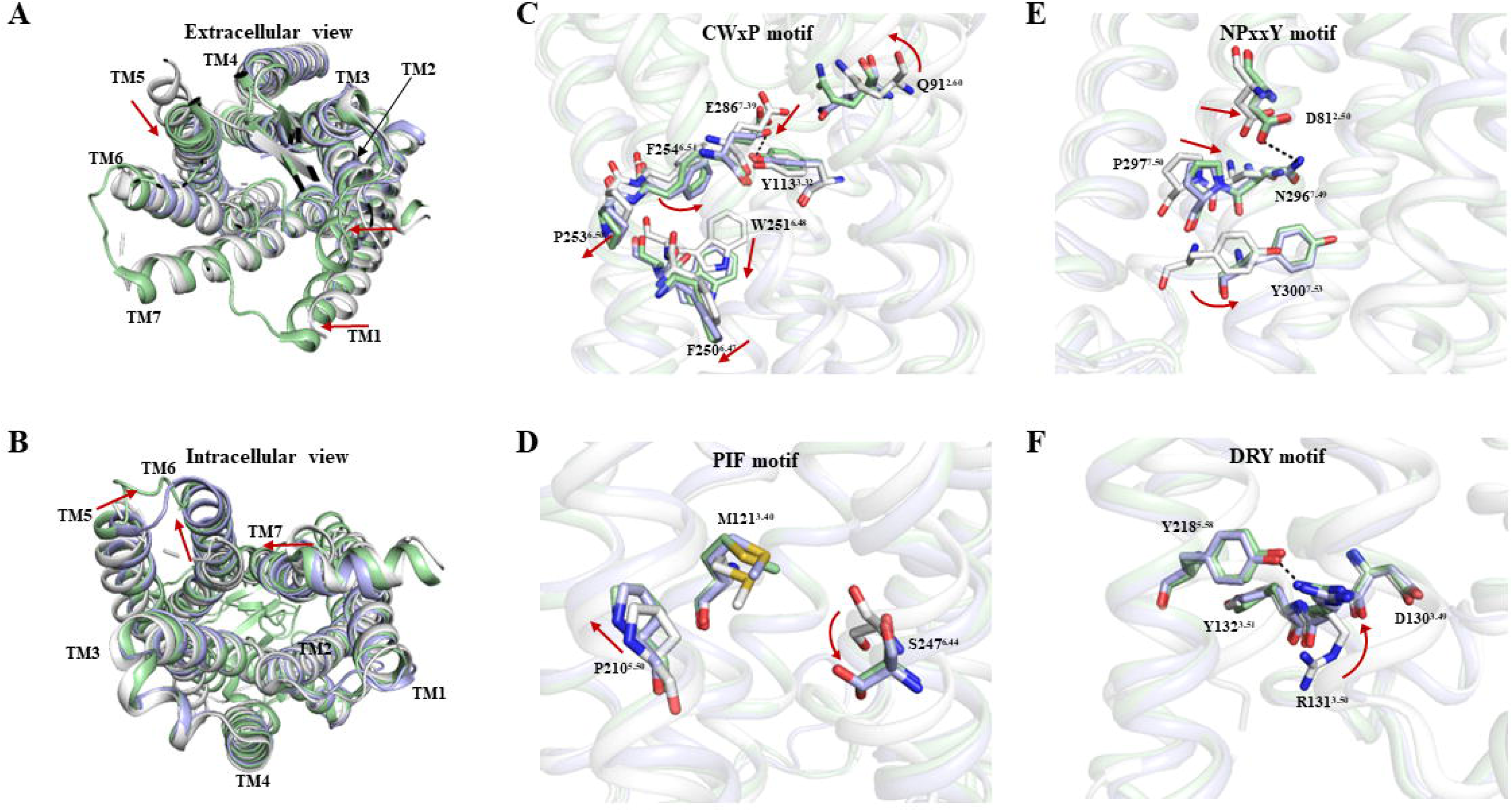
Molecular basis for the activation of CCR8. Comparison of CCR8-G_i_-scFv16 structure (light blue) with the structures in Fab1 mediated inactive-state (PDB ID 8TLM, gray) and CCL1 mediated active-state (PDB ID 8U1U, pale green) of CCR8. The structural superposition is displayed in the extracellular view **(A)** and intracellular view **(B)**. **(C-F)** Close-up views of the conserved CWxP **(C)**, PIF **(D)**, NPxxY **(E)**, and DRY **(F)** motifs show conformational changes along the pathway of receptor activation. Hydrogen bonds are depicted as black dashed lines.

### Activation of CCR8

The residue Q91^2.60^ rotated towards TM3, occupying a position similar to the binding site of residue M26 of CCL1. This rotation pushed residue E286^7.39^ of CCR8 downwards to the cytoplasmic side, and established a hydrogen bond interaction with the side chain of Y113^3.32^. F254^6.51^ is positioned between E286^7.39^ and W251^6.48^. Consequently, the movement of E286^7.39^ led to a deflection of F254^6.51^, and then induced the side-chain rearrangement of W251^6.48^ by steric hindrance (Fig. 4C). The movement of W251^6.48^ further triggered a rearrangement of the conserved PIF motif (P210^5.50^, M121^3.40^, and S247^6.44^ in CCR8) (Fig. 4D). Subsequently, the activation signal propagated through the conserved NPxxY motif (N296^7.49^, P297^7.50^, and Y300^7.53^ in CCR8) (Fig. 4E) to the bottom DRY motif (D130^3.49^, R131^3.50^, and Y132^3.51^ in CCR8) (Fig. 4F), ultimately causing TM6 to move outward to accommodate G protein binding. Two pairs of hydrogen bond interactions were present (R131^3.50^-Y218^5.58^ and D81^2.50^-N296^7.49^) to stabilize the active state of CCR8 (Fig. 4E and 4F).

### Interface between CCR8 and G_i_

Upon superposition of our structure with the CCL1-CCR8-G_i_ (PDB ID 8U1U), the outward movement of TM6 is less pronounced in our structure (with TM6 exhibiting an inward movement of 1.2 Å). However, TM5 in our structure is closer to TM6, reducing the distance between them from 8.8 Å to 8.2 Å. Additionally, the αN helix at the N-terminus of the Gα subunit undergoes an upward shift of 4.3 Å as shown in Fig. 5A, where the Gβγ subunits and scFv16 are omitted.

**Fig. 5.**
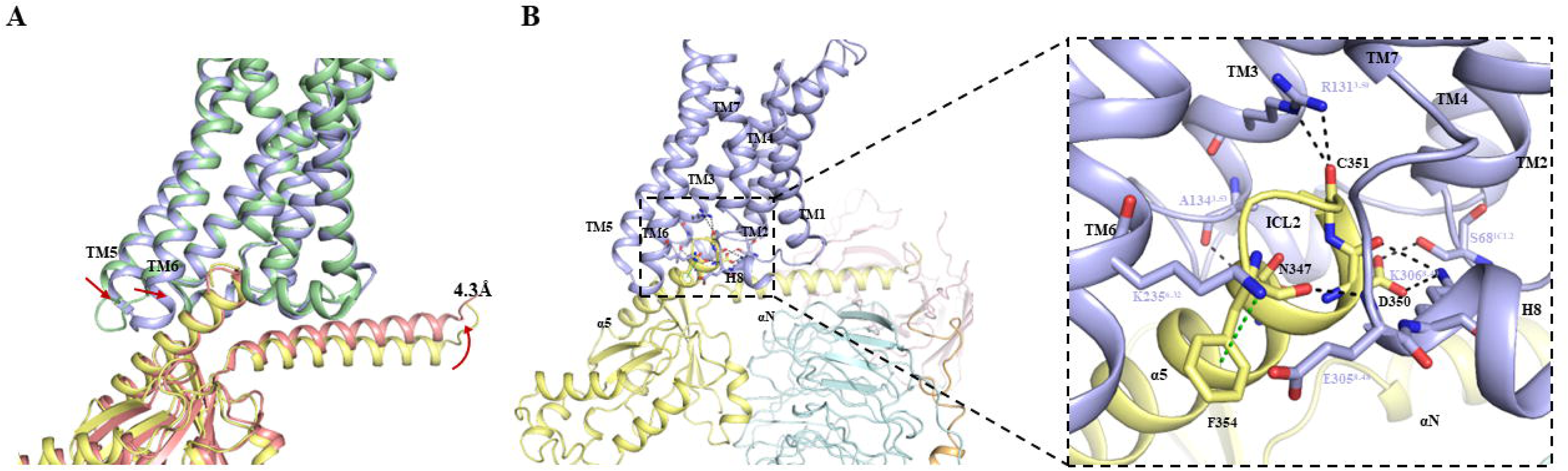
Interaction between CCR8 and G_i_. **(A)** Overview of CCR8-G_i_ interface. The structure of apo CCR8-G_i_ complex (CCR8 in light blue and Gα_i_ in pale yellow) is superposed with CCL1 activated CCR8-G_i_ structure (PDB ID 8U1U, CCR8 in pale green and Gα_i_ in salmon). TM6 exhibits a 1.2 Å inward movement in our apo CCR8-G_i_ structure compared to the CCL1-CCR8-G_i_ structure. TM5 is brought closer to TM6 in our apo CCR8-G_i_ structure, reducing the distance from 8.8 Å to 8.2 Å. The αN helix at the N-terminus of the Gα subunit shifts upward by 4.3 Å. **(B)** An enlarged view of **the** CCR8-G_i_ interface in our apo CCR8-G_i_ structure. Four hydrogen bond interactions (S68^ICL2^-D350^H5.22^, R131^3.50^-C351^H5.23^, A134^3.53^-N347^H5.19^, E305^8.48^-F354^H5.26^) and a salt bridge (K306^8.49^-D350^H5.22^) are observed and indicated as black dashed lines. A pi-cation interaction is formed between K235^6.32^ and F354^H5.26^ and is indicated as a green dashed line.

In general, the complex shares similarities in the interaction between the receptor and Gα_i_ protein with other class A GPCRs. The C-terminal α5-helix of Gα_i_, adopting a straight amphipathic α-helix, inserts into the cytoplasmic core of the transmembrane domain (TMD) and forms extensive hydrophobic interactions with TM3, intracellular loop 2 (ICL2), TM5, ICL3, and TM6 of the receptor. These interactions with the C-terminal α5-helix of Gα_i_ are primarily formed with the intracellular end of ICL2 and TM3, as well as the intracellular ends of ICL3, TM5, and TM6. Additionally, the residue S68^ICL1^ forms a hydrogen bond with D350^H5.22^. Residues located on TM3 directly interact with Gα_i_, including polar interactions such as R131^3.50^-C351^H5.23^ and A134^3.54^-N347 ^H5.19^. E305^8.48^ and K306^8.49^ are also involved in interactions with Gα_i_. Notably, K235^6.32^ forms a pi-cation interaction with F354 ^H5.26^ (Fig. 5B).

### CCR8-mediated calcium mobilization

The calcium mobilization assay is a cell-based second messenger technique used to quantify the calcium flux linked to the activation or inhibition of G-protein-coupled receptors. The alteration in fluorescence intensity is directly proportional to the quantity of intracellular calcium released into the cytoplasm in response to ligand activation of the target receptor. In this study, the calcium mobilization assay was employed to assess the effects of CCR8 residue mutations on the binding with a previously identified agonist, LMD-009. A total of 11 CCR8 mutants (Y42^1.39^A, F88^2.57^A, Q91^2.60^A, Q91^2.60^W, V109^3.28^A, S110^3.29^A, Y113^3.32^A, Y114^3.33^A, F254^6.51^A, L258^6.55^A, and E286^7.39^A) were included for LMD-009 binding investigation. These residues are likely involved in LMD-009 binding according to previous studies [20,25]. It was found that nine out of these 11 CCR8 mutants show little change in the EC_50_ values in LMD-009-induced calcium mobilization assay, compared with that of wildtype CCR8. However, two mutants, Y113A^3.32^ and E286A^7.39^, have significantly higher EC_50_ values (at least 3000-fold decrease) compared with that of wildtype CCR8, indicating the crucial importance of Y113^3.32^ and E286^7.39^ in the activation of CCR8 mediated by LMD-009 (Fig. 6). Notably, Y113^3.32^ and E286^7.39^ of CCR8 also exert an important role in the activation of CCR8 mediated by the endogenous ligand CCL1 [17]. These data provide implications for the ligand binding of CCR8.

**Fig. 6.**
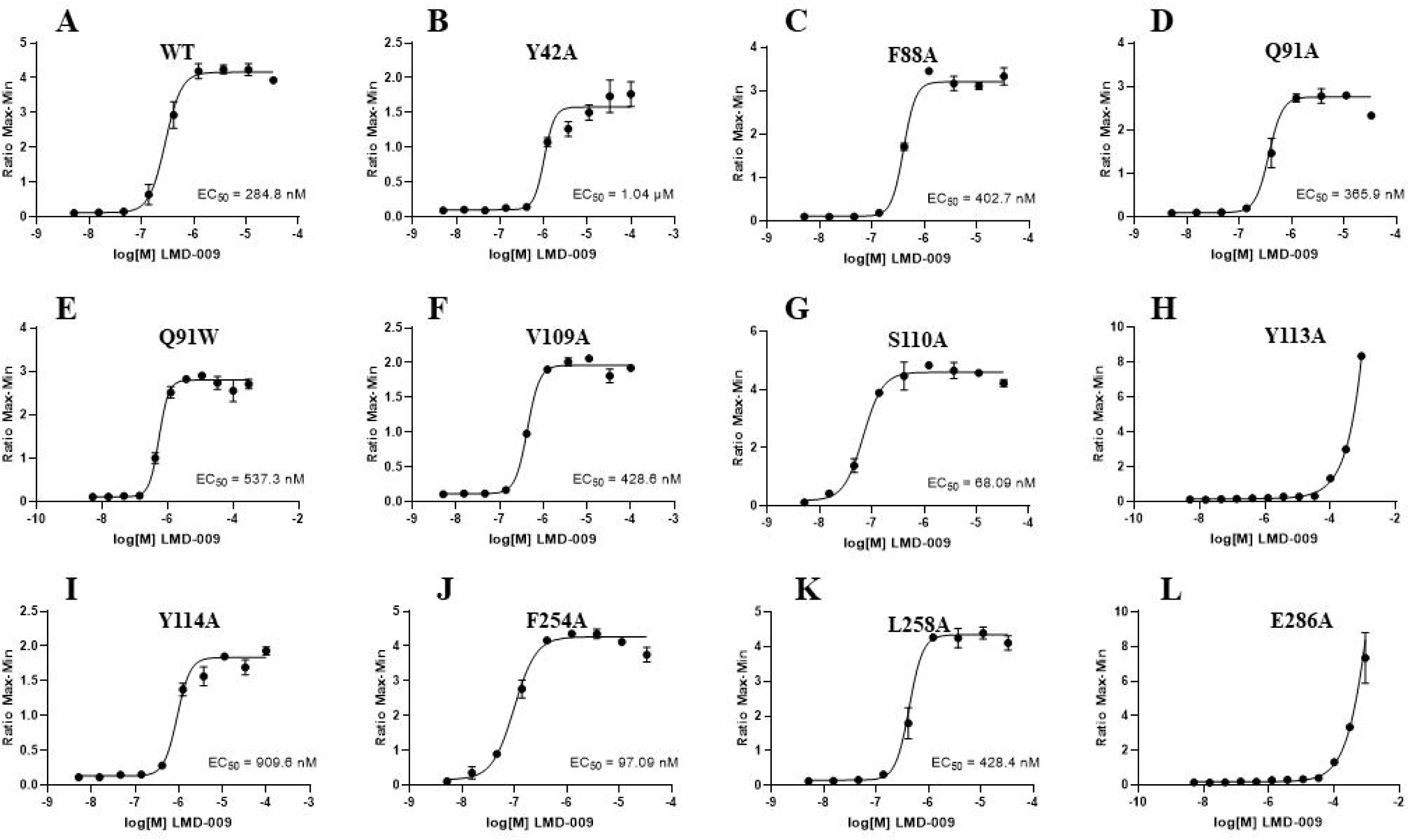
Effects of residue mutation of CCR8 on calcium mobilization induced by LMD-009. The calcium flux response of CCR8 induced by LMD-009 was detected by measuring the intracellular Ca^2+^ level. **(A)** Induced calcium mobilization of wildtype (WT) CCR8 by LMD-009. **(B-L)** The effects of residue mutations on LMD-009-induced activation of CCR8. Eleven mutations, namely Y42^1.39^A (B), F88^2.57^A (C), Q91^2.60^A (D), Q91^2.60^W (E), V109^3.28^A (F), S110^3.29^A (G), Y113^3.32^A (H), Y114^3.33^A (I), F254^6.51^A (J), L258^6.55^A (K), and E286^7.39^A (L) are included for investigation. Two mutations (Y113^3.32^ and E286^7.39^) of CCR8 result in significantly decreased calcium mobilization ability of LMD-009. The other mutations show little effect on calcium mobilization induced by LMD-009.

## Discussion

G-protein-coupled receptors (GPCRs), consisting of numerous structurally similar proteins, represent one of the most abundant protein classes encoded by the human genome. Located on the cell membrane and characterized by seven transmembrane helices, GPCRs are able to mediate signal transduction and take part in many essential physiological activities, thus being promising targets for drug development [1–4]. CCR8 is a member of the C-C motif chemokine receptor (CCR) subfamily of chemokine receptors. It has been a vital drug target in the treatment of cancer immunotherapy as well as autoimmune diseases [3–7]. Although the structures of human CCR8 in complex with an antagonist antibody (Fab1) and an endogenous agonist ligand CCL1 have been determined [17], the structure of CCR8 in apo state remains to be solved. Limited structural information on human CCR8 will hamper the generation of therapeutics against this emerging target. In the present study, we solved the cryo-EM structure of CCR8 in a complex with G proteins. This is the first report of the cryo-EM structure of ligand-free CCR8. Together with previous structural studies of CCR8 [17], this study will enrich the structural information of CCR8 and, absolutely, will add to the development of CCR8-targeted therapy.

Compared to the structures of the chemokine-CCR8-G_i_ complexes and inactive CCR8, we found that CCR8 in our solved ligand-free CCR8-G_i_ structure displayed a typical active state. Increasing evidence supports the self-activation of GPCRs [26–29], while this study firstly reported the active features of CCR8 in apo structure, which is due to some conformational changes of residues in the ligand binding pockets. These observations will expand our knowledge of the diversity of GPCR activation.

Agonists targeting CCR8 hold great potential to treat autoimmune diseases. LMD-009 is a robust nonpeptide agonist of CCR8, and molecular interaction between LMD-009 and CCR8 was investigated by mutagenesis of CCR8 and calcium mobilization assays previously [20]. This study provided similar but more data about CCR8-LMD-009 interaction. Both the present study and the previous report demonstrated that mutations of residues Y113^3.32^ and E286^7.39^ of CCR8 significantly reduce the LMD-009 mediated calcium mobilization level. The structure of CCR8 in complex with LMD-009 is expected to be solved to demonstrate the strong interaction between Y113^3.32^ and E286^7.39^ of CCR8 with LMD-009.

## Materials and Methods

### Construct design, Protein Expression and purification

The human CCR8 gene (UniProtKB-P51685) was cloned into a modified pFastBac1 vector with an N-terminal hemagglutinin (HA) signal sequence, a Flag-tag (DYKDDDD), a thermostabilized apocytochrome b562RIL (BRIL) and a C-terminal tobacco etch virus (TEV) protease cleavage site, followed by a 10×His tag. A mutation F122^3.41^W was introduced to enhance protein yield and homogeneity. For the assembly of the CCR8-G_i_ complex, a NanoBiT tethering method was employed. This involved fusing the C-terminus of CCR8 to the large part of NanoBiT (LgBiT) and attaching the C-terminus of Gβ_1_ to the small part of NanoBiT (HiBiT). Human Gα_i1_ and Gβ_1_γ_2_ subunits were cloned into pFastbac1 and pFastBac Dual vectors, respectively. To reduce nucleotide binding and increase the stability of the Gα_i1_β_1_γ_2_ complex, we employed a dominant-negative Gα_i1_ (DNGα_i1_) format, which included mutations of S47C, G202T, G203A, E245A, and A326S. Additionally, the scFv16 gene was integrated into the pFastBac1 vector, incorporating an N-terminal GP67 signaling peptide and a C-terminal HRV3C protease cleavage site, followed by an 8× His-tag. CCR8, DNGα_i1_, Gβ_1_γ_2_ and scFv16 were co-expressed in Spodoptera frugiperda (Sf9) with the addition of all four baculoviruses (ratio of 1:2:1:1 (CCR8:DNGα_i1_:Gβ_1_γ_2_:scFv16)) to Sf9 cells at a density of 1.5×10^6^ cells/mL. 1 μM of LMD-009 (MedChemExpress) was added to the media during growth. After 48 h, cells were collected and stored at −80 °C. When used, sf9 cell pellets were resuspended and lysed in a hypotonic buffer consisting of 20 mM HEPES (pH 7.5), 100 mM NaCl, 10 mM MgCl_2_, 20 mM KCl, 5 mM CaCl_2_, 100 μM TCEP, 5 μM LMD-009 and EDTA-free protease inhibitor cocktail tablets (Roche). Then the suspension was incubated at room temperature for 1.5 h after adding apyrase (25 mU/mL; New England Biolabs). The supernatant was removed by ultracentrifugation at 40,000 rpm for 30 min, and the membrane pellet was resuspended and solubilized in a buffer containing 20 mM HEPES (pH 7.5), 100 mM NaCl, 5 mM CaCl_2_, 100 μM TCEP, 5% glycerol, 5 μM LMD-009, EDTA-free protease inhibitor, apyrase (50 mU/mL). Subsequently, 0.5% (w/v) lauryl maltose neopentyl glycol (LMNG, Anatrace) and 0.05% (w/v) cholesteryl hemisuccinate TRIS salt (CHS, Anatrace) were added, and solubilization proceeded for 2 h at 4 °C, followed by centrifugation at 40,000 rpm for 40 min at 4 °C. The supernatant was collected and incubated with TALON IMAC resin (Clontech) at 4 °C for 2 h in the presence of 20 mM imidazole. The resin was packed onto a disposable gravity column (Bio-Rad) and washed with 10 column volumes (CV) of Wash Buffer I (20 mM HEPES, pH 7.5, 100 mM NaCl, 5% glycerol, 0.1% (w/v) LMNG, 0.01% (w/v) CHS, 20 mM imidazole, 100 μM TCEP, 5 μM LMD-009), followed by 10 CV of Wash Buffer II (20 mM HEPES, pH 7.5, 100 mM NaCl, 5% glycerol, 0.05% (w/v) LMNG, 0.005% (w/v) CHS, 30 mM imidazole, 100 μM TCEP, 5 μM LMD-009). The complex protein was then eluted by 3CV of an elution buffer containing 20 mM HEPES (pH 7.5), 100 mM NaCl, 5% glycerol, 0.03% (w/v) LMNG, 0.003% (w/v) CHS, 300 mM imidazole, 100 μM TCEP, 5 μM LMD-009. The eluate was concentrated and applied onto a Superose 6 Increase 10/300 column (GE Healthcare) for size exclusion chromatography (SEC) with buffer containing 20 mM HEPES, pH 7.5, 100 mM NaCl, 0.003% (w/v) LMNG, 0.0003% (w/v) CHS, 100 μM TCEP, 1 μM LMD-009. Fractions were pooled together and concentrated using a 100 kDa molecular mass cut-off concentrator (Millipore) for cryo-EM grid preparation.

### Cryo-EM sample preparation and data acquisition

To prepare the cryo-EM grid of the CCR8-G_i_-scFv16 complex, 3μL of the purified samples at 6.2 mg/mL were added to 300 Mesh R1.2/1.3 Au Quantifoil grids (glow discharged at 15mA for 40 seconds with a Glow discharge cleaning system). Grids were blotted with qualitative filter paper in a Vitrobot Mark L (Thermo Fisher Scientific) at 4L and 100% humidity for 3 seconds and plunge frozen in liquid ethane, and then stored in liquid nitrogen until checked. We used the 300LkV Titan Krios Gi3 microscope (Thermo Fisher Scientific FEI, the Kobilka Cryo-EM Center of the Chinese University of Hong Kong, Shenzhen) to check the grids and collect the cryo-EM data of the CCR8-G_i_-scFv16 complex. The Gatan K3 BioQuantum camera at a magnification of 105,000 was used to record movies, and the pixel size was 0.83 Å. The movie stacks were automatically acquired with the defocus range from −1.0 to −2.0 μm. The exposure time was 2.5Ls, with frames collected for a total of 50 frames (0.05 s/frame) per sample with a total dose of 45 e^−^/Å^2^. SerialEM 3.7 was used for semiautomatic data acquisition.

### Cryo-EM data processing

For the CCR8-G_i_-scFv16 complex, a total of 3996 movies stacks were imported into cryoSPARC v4.1.1. After motion corrected, electron-dose weighted and CTF estimation filtered out movies with a resolution greater than 4 Å. The initial particle was performed by cryoSPARC blob picker. A total number of 2,694,600 particles were extracted with the blob picking. After four rounds of 2D classification, the number of good particles was reduced to 926,335. The number of particles was further reduced to 277,386 by heterogeneous refinement and Ab-initio reconstruction. The nonuniform refinement enables us to reconstruct the 3.43 Å structure. To further improve the resolution, the final particle sets were re-extracted with the original box size and further applied for final nonuniform refinement and local refinement in cryoSPARC. A density map was obtained with an overall resolution of 2.58 Å (determined by gold standard Fourier shell correlation (FSC) using the 0.143 criterion).

### Model building and refinement for cryo-EM structures

Human CCR8 maps were visualized in UCSF Chimera. The Cryo-EM structure of the CCR5-G_i_ complex (PDB ID 7O7F) was used as a reference to build the initial model. After that, the models were subjected to several rounds of manual adjustment and auto-refinement in Coot and Phenix, respectively [30,31]. The quality of the final model was validated by MolProbity [32]. The specific parameters for cryo-EM data collection, refinement, and validation statistics are presented in Table S1. Structure figures were prepared using PyMOL and Chimera.

### Calcium mobilization assay

Calcium mobilization assay is a cell-based second messenger assay to measure the calcium flux associated with G-protein coupled receptor activation or inhibition. The change in the fluorescence intensity is directly correlated to the amount of intracellular calcium that is released into the cytoplasm in response to LMD-009 activation of the receptor of interest. This assay is employed to assess the impact of amino acid mutations in the CCR8 receptor on CCR8 activation. CCR8 (WT and mutants) were cloned into the pLVX-Puro vector. The CCR8-Gqi5-CHO-K1 cells (constructed by Genomeditech) passed in a complete medium (1640, 10% FBS, 1 μg/mL puromycin) in an incubator (37L, 5% CO_2_) were used in the Calcium mobilization assay.

The fluorescent membrane-permeable calcium-binding dye (the FLIPR Calcium 6 Assay Kit) was dissolved in assay buffer (20 mM HEPES buffer with 1×Hank’s Balanced Salt Solution (HBSS), pH 7.4). The loading buffer was prepared with the dye solution containing 5 mM probenecid. The probenecid was prepared into 500 mM stock solution in 1 N NaOH, and then diluted to 250 mM in HBSS buffer before use.

Approximately 1.5×10^4^ CCR8-Gqi5-CHO-K1 cells were seeded into a 384-well plate and incubated in 25 μL complete medium 1640, 10% FBS, 1 μg/mL puromycin) in 5% CO_2_ at 37L for 16 hours. Then, the complete medium was completely changed with 25 μL assay buffer, and 25 μL loading buffer was added to the desired wells. After adding dye, the cell plate was incubated for 2 hours at 37°C with 5% CO_2_ and then kept at room temperature until used. The microplate was transferred to the FLIPR instrument, and the calcium assay was started as described in the user guide for the instrument. 12.5 μL assay buffer containing LMD-009 with a concentration gradient was added during the assay. The MAX-MIN ratio value was plotted against the LMD-009 concentration and analyzed in GraphPad Prism for concentration curve generation.

## Data deposition

The cryo-EM electron density map of human CCR8 has been deposited in the Electron Microscopy Data Bank (accession number EMD-38481), and the fitted coordinate has been deposited in the Protein Data Bank (PDB ID 8XML).

## Acknowledgments

J.L. was supported by Jiangxi Natural Science Foundation for Distinguished Young Scholars (20212ACB216001), Jiangxi Key Research and Development Program (20203BBG73063), Jiangxi ‘‘Double Thousand Plan’’ (jxsq2019101 064), and the foundation of Gannan Medical University (QD201910). J.Z. was supported by the National Natural Science Foundation of China (32271260), the CAS ‘‘Light of West China’’ Program (xbzg-zdsys-202005), and the Natural Science Foundation of Jiangxi Province (grant number: 20224ACB206046). H.J. was supported by the National Natural Science Foundation of China (32360223).

## Author Contributions

Q.P. performed the experiments (including designing and generating recombinant DNA plasmids, optimized expression and purification of CCR8-G_i_-scFv16), interpreted the data, prepared the figures, and wrote the first draft of the manuscript (ms); H.J. contributed to the writing and revision of the ms; X.C. carried out cryo-EM data processing; N.W. prepared mutants and performed the calcium mobilization assay; W.N. contributed to the expression of Sf9 cells and the harvesting cells; G.F. supervised calcium mobilization assay; J.L. contributed to building the atomic model on the basis of cryo-EM maps and revised the ms; J.Z. initiated the project, conceived the study, designed the experiments, interpreted the data, wrote the ms and supervised for the research. The other authors all coordinated the study.

