## Supplemental Table1 for "Cryo-EM structure and biochemical analysis of human chemokine receptor CCR8"

**Table S1.** Cryo-EM data collection, refinement and validation statistics

|  | CCR8-G_i_-scFv16  (EMD-38481)  (PDB 8XML) |
| --- | --- |
| **Data collection and processing** |  |
| Magnification | 105,000 |
| Voltage (kV) | 300 |
| Electron exposure (e–/Å^2^) | 45 |
| Defocus range (μm) | -1.0−-2.0 |
| Pixel size (Å) | 0.83 |
| Symmetry imposed | C1 |
| Initial particle images (#) | 2,694,600 |
| Final particle images (#) | 275,730 |
| Map resolution (Å)  FSC threshold | 2.58  0.143 |
| **Refinement** |  |
| Initial model used (PDB code) | 7O7F |
| Model resolution (Å)  FSC threshold | 3.15  0.143 |
| Model composition  Non-hydrogen atoms  Protein residues  Ligands | 8237  1049  0 |
| r.m.s. deviations  Bond lengths (Å)  Bond angles (°) | 0.01  0.79 |
| Validation  MolProbity score | 1.46 |
| Ramachandran plot  Favored (%)  Allowed (%)  Disallowed (%) | 96.09%  3.91%  0 |
